# Agonists of orally expressed TRP channels stimulate salivary secretion and modify the salivary proteome

**DOI:** 10.1101/2020.06.17.157198

**Authors:** Jack William Houghton, Guy Carpenter, Joachim Hans, Manuel Pesaro, Steven Lynham, Gordon Proctor

## Abstract

Natural compounds that can stimulate salivary secretion are of interest in developing treatments for xerostomia, the perception of a dry mouth, that affects between 10 and 30% of the adult and elderly population. Chemesthetic transient receptor potential (TRP) channels are expressed in the surface of the oral mucosa. The TRPV1 agonists capsaicin and piperine have been shown to increase salivary flow when introduced into the oral cavity but the sialogogic properties of other TRP channel agonists have not been investigated. In this study we have determined the influence of different TRP channel agonists on the flow and protein composition of saliva.

Mouth rinsing with the TRPV1 agonist nonivamide or menthol, a TRPM8 agonist, increased whole mouth saliva (WMS) flow and total protein secretion compared to unstimulated saliva, the vehicle control mouth rinse or cinnamaldehyde, a TRPA1 agonist. Nonivamide also increased the flow of labial minor gland saliva but parotid saliva flow rate was not increased. The influence of TRP channel agonists on the composition and function of the salivary proteome was investigated using a multi-batch quantitative mass spectrometry method novel to salivary proteomics. Inter-personal and inter-mouth rinse variation was observed in the secreted proteomes and, using a novel bioinformatics method, inter-day variation was identified with some of the mouth rinses. Significant changes in specific salivary proteins were identified after all mouth rinses. In the case of nonivamide, these changes were attributed to functional shifts in the WMS secreted, primarily the over representation of salivary and non-salivary cystatins which was confirmed by immunoassay.

This study provides new evidence of the impact of TRP channel agonists on the salivary proteome and the stimulation of salivary secretion by a TRPM8 channel agonist, which suggests that TRP channel agonists are potential candidates for developing treatments for sufferers of xerostomia.

## Introduction

TRP (Transient Receptor Potential) channels are a superfamily of non-selective cation channels that respond to a variety of somatosensory and endogenous stimuli. TRPV1, 3, 4, TRPA1 and TRPM8 are expressed in the oral cavity that have thermo- and chemoreceptive functions. They are expressed on mucosal and epithelial free afferent nerve endings of myelinated Aδ and non-myelinated C fibres (1), oral epithelial cells (2-4), taste buds (5, 6), and keratinocytes (7).

Cinnamaldehyde is a TRPA1 agonist, which is produced synthetically and found in cinnamon, a spice that comes from the bark of cinnamon trees (8). Cinnamaldehyde makes up 90% of the essential oil extracted from cinnamon bark. Upon contact, cinnamaldehyde provokes a feeling of warmth (8) and has potential anti-inflammatory (9-11) and anti-cancer (12-18) properties. Menthol is a TRPM8 agonist that provokes a cooling sensation. It is found in mint leaves and produced synthetically (19). Nonivamide is a capsaicinoid that elicits a burning sensation (20). It is structurally very similar to the more widely studied TRPV1 agonist capsaicin and is naturally found in chilli peppers or produced synthetically.

The salivary response to basic tastants is well studied but the salivary response to TRP channel agonists requires further investigation. Increased salivary flow rate and specific protein secretion have been demonstrated in response to other tastants (21-24) and there are studies demonstrating increases in salivary flow rates and specific protein changes in response to the TRPV1 agonists (25-29) but there has been limited study of agonists to other TRP channels, despite expression of these channels in the oral cavity, nor has the mechanism of TRP channel agonist stimulated salivary secretion been elucidated.

Studying compounds that can stimulate salivary flow is of interest to the development of treatments for xerostomia, the perception of a dry mouth, that affects between 10 and 30% of the adult and elderly populations (30, 31). Acidic tastants that strongly stimulate salivary secretion erode enamel tissues, so alternative molecules are sought (32). Although xerostomia is often associated with hyposalivation, where the WMS flow rate is reduced by ∼50% (33), this is not always the case (34). Xerostomia in the absence of hyposalivation may be due to changes in the interaction of saliva with oral surfaces due to the altered integrity of salivary proteins (35) or changes in saliva rheology (36). There is evidence that TRP agonists modify the rheological properties of saliva but the mechanism by which these changes occur remains to be elucidated. Taken together, identifying compounds that not only induce salivary secretion but also modify the rheological properties of saliva is of interest to developing treatments for xerostomia.

Specific protein changes in saliva in response to differing stimuli are possible due to the many sources of proteins which are likely to respond differently to different nerve mediated stimuli. For example, the submandibular and sublingual glands secrete in response to olfaction (37) whereas the parotid glands do not (38). Conversely, the parotid glands are preferentially stimulated by chewing which results in a higher amylase output (39). In these scenarios, proteins associated with specific glands, e.g. higher amylase secretion by the parotid glands or mucin secretion by the submandibular and sublingual glands, will have a relatively increased abundance when compared to unstimulated levels.

The regulation of specific proteins separate from preferential gland stimulation has also been reported. Annexin A1 and calgranulin A are upregulated in WMS through an inflammatory-like response after mouth rinsing with bitter, umami and sour tastants (40). Bader et al. demonstrated the upregulation of lysozyme in saliva stimulated by citric acid rinse (41). The TRPV1 agonist 6-gingerol upregulated salivary sulfhydryl oxidase 1 resulting in reduced 2-furfurylthiol levels in exhaled breath and thus reduction in the perceived sulphur-like after-smell (42). However, the mechanism of these specific protein upregulations has not been elucidated.

The present study is formed of two parts. A bottom-up quantitative proteomics study of the salivas secreted by two participants in response to menthol, cinnamaldehyde, nonivamide and propylene glycol (PG) that were compared to unstimulated saliva using mass spectrometry. In addition, data on WMS flow rates and protein output were also collected. In order to improve the identification of lower abundance salivary proteins, a method novel to salivary proteomics was used. Secondly, studies were conducted to confirm the specific protein changes of the proteomes of salivas identified in the proteomics study and to consider the mechanism by which the compounds exert their effects on the salivary proteomes.

## Experimental Procedures

### Experimental Design and Statistical Rationale

For the proteomics study, the proteome of 60 WMS samples, obtained from two male volunteers of ages 24 and 27, were analysed by TMT quantitative mass spectrometry. Forty eight experimental samples consisting of WMS produced after mouth rinsing were split randomly across six TMT10plex batches with each batch containing two controls consisting of pooled unstimulated saliva from each participant. The 48 WMS samples were collected from two participants after being exposed to eight conditions each with three experimental repeats. In a further study of the effects of agonists on WMS secretion, 25 participants were recruited (the demographic information of each participant group is shown in Table 1) six of these subjects also participated with further participants in the following studies. For the parotid saliva study, eight volunteers were recruited (38.7 ± 5.3 years, male *n* = 4, female *n* = 4). For the lower labial gland saliva study, ten volunteers were recruited (29.4 ± 4.7 years, male *n* = 5, female *n* = 5). For all studies, volunteers were healthy individuals recruited by internal advertisement with the following exclusion criteria: on prescription medication, age > 65years or < 18years, suffering from oral discomfort. The controls and statistical tests used for each analysis are described below.

**Table 1.** Demographic information of participants in the WMS study.

| Study | Mean age | SEM | *n* | Male | Female |
| --- | --- | --- | --- | --- | --- |
| Nonivamide | 25.3 | 2.1 | 7 | 4 | 3 |
| Menthol | 27.2 | 1.5 | 6 | 3 | 3 |
| Cinnamaldehyde | 27.6 | 4.1 | 6 | 3 | 3 |
| PG | 27.2 | 2.5 | 6 | 3 | 3 |

#### Proteomics study of TRP agonist stimulation on two subjects

Forty eight saliva collections were made in total, each collection including an unstimulated saliva sample, followed by a mouth rinse and then two post-mouth rinse saliva samples (Table 2). Eight different mouth rinse solutions were tested in triplicate: nonivamide, cinnamaldehyde, menthol and PG (Symrise AG) (Table 2). The solutions were prepared in pre-weighed universal tubes and the total weight recorded. The compounds were diluted in water (Buxton) on the day of collection and were stored at room temperature. Participants were asked not to consume food, water or smoke in the 1 hour prior to collection. The following guidance was given to each participant prior to each collection: tilt your head slightly forward to allow saliva to pool underneath the tongue; do not move your mouth unless it is to spit out collected saliva; spit out whenever it is comfortable; do not swallow. For each collection, the following protocol was adhered to: One minute of unstimulated WMS was collected in a pre-weighed universal tube; 10 mL of mouth rinse was then taken into the mouth for 30 seconds and spit back into a pre-weighed universal tube; two, one minute collections of post-mouth rinse WMS in pre-weighed universal tubes. Immediately after collection, participants were asked, “How would you rate the intensity of the mouth rinse” and were asked to give a rating from 0 – 10 on a visual analogue scale alongside an oral description of their perception of the mouth rinse. One collection was carried out per day at 2pm and the order of mouth rinses were randomised for each participant. All samples were weighed in the universal tube straight after collection. Saliva was then processed for storage prior to mass spectrometry analysis: samples were transferred to ice cooled 1.5 mL microtube for centrifugation (13 500 rpm, 5 minutes, 4 °C). Supernatants were removed, frozen at -20 °C and finally moved to -80 °C storage; the pellets were discarded.

**Table 2.** The concentrations of mouth rinses used in each saliva collection of the proteomics study. Each collection consisted of an unstimulated saliva sample, followed by a 30 second mouth rinse and then 2 × 1 minute post-mouth rinse saliva samples. Each collection was carried out in triplicate for two participants, totalling 48 collections. The compound, concentration and PG content in each of the mouth rinses used for this study are shown in the table.

| Compound | Concentration (ppm) | PG dilution |
| --- | --- | --- |
| PG | $1.8 \times 10^4$ | n/a |
| PG | $3.0 \times 10^4$ | n/a |
| Menthol | 300 | $6.0 \times 10^3$ ppm |
| Menthol | 500 | $1.0 \times 10^4$ ppm |
| Cinnamaldehyde | 180 | $1.8 \times 10^4$ ppm |
| Cinnamaldehyde | 300 | $3.0 \times 10^4$ ppm |
| Nonivamide | 0.6 | $6.0 \times 10^2$ ppm |
| Nonivamide | 1.0 | $1.0 \times 10^3$ ppm |

### WMS Saliva Collection

#### Effects of TRP agonists on WMS flow rates

Cinnamaldehyde, menthol and nonivamide were obtained from Symrise AG and prepared in PG. 300 ppm cinnamaldehyde, 500 ppm menthol, 1ppm nonivamide and 3 × 10^4^ ppm PG were prepared by diluting in water (Buxton) in pre-weighed universal tubes and the total weights were recorded. The concentration of PG in the nonivamide, menthol and cinnamaldehyde mouth rinses was 3 × 10^3^, 1 × 10^4^ and 3 × 10^4^ ppm respectively. The solutions were kept at room temperature (20 °C). Participants were asked not to consume food, water or smoke in the 1 hour prior to collection. Prior to collection each participant was asked to tilt their head slightly forward to allow saliva to pool underneath the tongue, to not move their mouth unless it was to spit out collected saliva, to spit out whenever it is comfortable and to not swallow. Five minutes of unstimulated WMS was collected in a pre-weighed universal tube as a control. Ten mL of a control mouth rinse containing either the equivalent concentration of PG as in the TRP agonist containing mouth rinse or water was then taken into the mouth for 30 seconds and spat back into a pre-weighed universal tube, this was followed by five one minute collections of WMS into pre-weighed universal tubes. This was repeated with the experimental mouth rinse. All samples were weighed in the universal tube immediately after collection. Samples were kept on ice after collection. The neat saliva samples were aliquoted into 2 mL microtubes and then centrifuged (13 500 rpm, 4 °C, 5 minutes). The supernatant was removed, aliquoted and stored at -20 °C.

### Parotid Saliva Collection

Five 10 mL solutions were prepared: water (Buxton); propylene glycol (3.0 × 10^4^ ppm), menthol (100 ppm), cinnamaldehyde (60 ppm), nonivamide (1 ppm). These solutions were prepared in pre-weighed universal tubes and the total weights recorded. The solutions were kept at room temperature (20 °C). Lashley cups were fitted over the exit of the Stenson’s ducts, secured and correct fitting was tested by the administration of a few drops of 2% citric acid onto the tongue to stimulate parotid secretion. Time was allowed so that the collection tubes of the Lashley tubes were filled with parotid saliva. Prior to collection each participant was asked to not swish any solution around in their mouth in order to prevent Lashley cups being dislodged. The volunteer was given 10 mL water to practice holding the solution in the mouth and spitting it out. Unstimulated parotid saliva was collected in a pre-weighed universal tube for 5 minutes. Ten mL of water (Buxton) was then taken into the mouth and held for 5 minutes. During this time parotid saliva was collected in a pre-weighed universal tube. This was repeated with the control and TRP agonist solutions in the following order: propylene glycol, menthol, cinnamaldehyde, and nonivamide. A two minute break was taken between each solution. Saliva samples were kept on ice after collection. The neat saliva samples were aliquoted into 2 mL microtubes and then centrifuged (13 500 rpm, 4 °C, 5 minutes). The supernatant was removed, aliquoted and stored at -20 °C.

### Lower labial gland saliva collection

A cotton roll was placed over each Stenson duct’s papilla and under the tongue to absorb major gland saliva. The inferior labial surface was dried, and unstimulated lower labial saliva was allowed to bead on the surface of the inferior labium for 2 minutes. A 2 cm x 1 cm piece of pre-weighed Whatman’s (General Electric) filter paper was then placed on the lower labial surface with one of the 1 cm edges halfway down the mid-point of the inferior labium to collect the beads of saliva. The saliva-soaked filter paper was placed in a pre-weighed 1.5 mL microtube, weighed and the flow rate calculated by subtraction of the pre-weighed paper and pre-weighed microtube weights and divided by the time of collection in minutes. To allow for slight variations in the size of the filter paper, flow rates were scaled according to the mass of the dried filter paper. This process was repeated but with a 30 second mouth rinse of either 3.0 × 10^4^ ppm PG, 300 ppm cinnamaldehyde, 500 ppm menthol or 1 ppm nonivamide being administered prior to the drying of the inferior labium. The following guidance was given to each participant prior to collection: ensure the mouth rinse baths the surface of your lower lip; do not swallow the mouth rinse. A three minute break, or until the perception of the previous mouth rinse had diminished, was taken between each solution. Saliva infused filter paper samples were kept on ice after collection.

Saliva infused filter paper was placed into 0.5 mL microtubes that had 4 needle-sized holes pierced into their underside. Each 0.5 mL microtube was then placed into a 1.5 mL microtube and centrifuged (13 000 rpm, 4 °C, 5 minutes). The saliva collected in the 1.5 mL microtube was immediately processed for SDS PAGE (see below) with the following modification: the entire volume of the collected saliva (∼1 µL) was treated with 10 µL lithium dodecyl sulphate (LDS) sample buffer and 1 µL dithiothreitol (DTT) prior to heating and electrophoresis.

### Quantitative tandem mass spectrometry

The first minute and second minute post-mouth rinse samples from each collection were pooled. The 24 unstimulated samples from each of the two participants (48 in total) were pooled into two unstimulated pools, one for each participant. Five µL of each pooled sample was added to 95 µL phosphate buffered saline (137mM NaCl, 2.7 mM KCl, 10 mM Na_2_HPO_4_, 1.8 mM KH_2_PO_4_, pH 7.4) for protein quantification using a Bradford assay (Thermo Scientific, USA). Absorbance of each sample was read by spectrophotometer at 595 nm and compared to a standard curve of bovine serum albumin of known protein concentration. Fifty µg of protein was extracted from each sample and frozen at -80°C. Frozen samples were freeze dried and reconstituted in 70 µL 100 mM triethylammonium bicarbonate (TEAB) and 0.1% sodium dodecyl sulphate (SDS). 10 µL 8mM tris (2-carboxyethyl) phosphine (TCEP) in 100 mM TEAB, 0.1% SDS was added to each sample and incubated at 55°C for one hour. 10 µL 375 mM iodoacetamide (IAA) in 100 mM TEAB, 0.1% SDS was added to each sample and incubated at room temperature for 30 minutes. 4 µL of 0.25 µg/µL trypsin (Roche, sequencing grade) was added to each sample and left overnight at 37 °C.

Forty one µL of TMT reagent was added to each of the 48 post mouth rinse samples and the twelve unstimulated pool samples (see Table 3 for details) and incubated at room temperature for one hour. Eight µL of 5% hydroxylamine was added to each sample and left at room temperature for 15 minutes. Samples from each 10plex batch were pooled into si× 10plex sample pools and stored at -80 °C prior to freeze drying until completion.

**Table 3.**
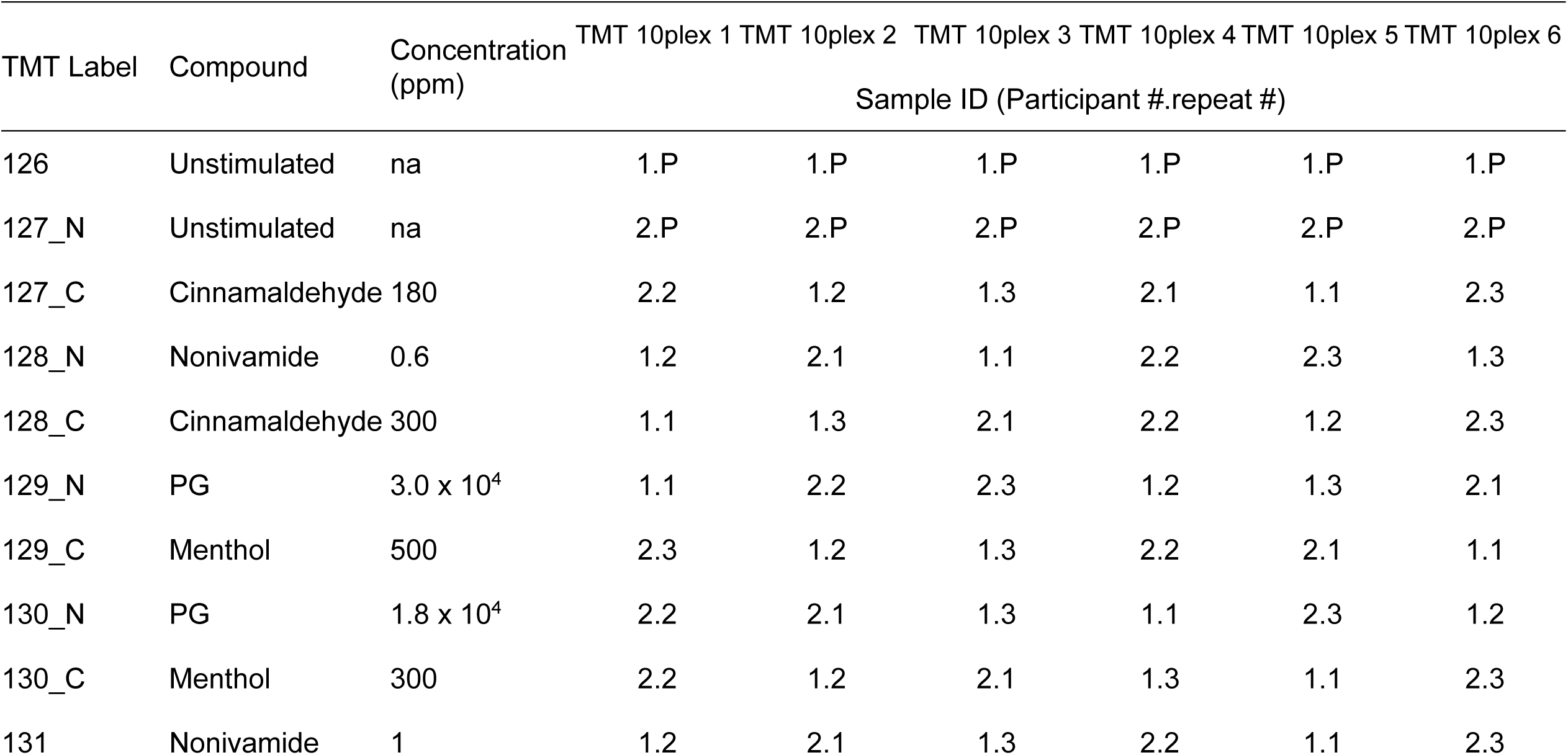
Quantitative analysis of the salivary proteome: TMT 10plex batch information. Note: P=pool.

IEF fractionation was carried out using the Agilent 3100 OFFGEL system (Agilent Technologies Inc, Germany) and was carried out according to the manufacturers protocol. 1.8 mL OFFGEL buffer stock added to each sample for reconstitution. Six OFFGEL strips with a linear pH gradient ranging from 3 to 10, one for each 10plex sample pool, were hydrated in 50 µL OFFEGL rehydration solution for 15 minutes. 12-fraction frames were fitted to each of the strips and 150 µL of reconstituted sample loaded into each fraction well. IEF was carried out under the following conditions: 20 kVh (100 hours, V: 500-5400 V, max. I: 50 µA. Upon completion, each fraction was removed and frozen at -80 °C. Fractions were thawed on ice and pooled into six fraction pools (Fraction 1 with 7, 2 with 8, 3 with 9, 4 with 10, 5 with 11 and 6 with 12). Ten µL of elution buffer (50% acetonitrile (ACN), 0.1% formic acid) was added to each sample. Zip-Tips were hydrated twice in 10 µL hydration solution (50% ACN, trifluoroacetic acid (TFA)) and then washed in 1 µL of wash solution (0.1% TFA). S10 µL samples was washed through the Zip-Tip 10 times before eluting with elution solution (0.1% TFA). The elute was frozen at -80 °C prior to freeze drying until completion. Fractions were reconstituted in 10 µL 50mM ammonium bicarbonate. The peptides from each fraction were resolved using reverse-phase chromatography on a 75 µM C18 EASY column using a 3-step gradient of 5-40% ACN and a 95% ACN wash in 0.1% formic acid at a rate of 300 µL/min over 220 minutes (EASY-NanoLC, ThermoScientific, USA). Nano-ESI was performed directly from the column and ions were analysed by using an LTQ Orbitrap Velos Pro (ThermoScientific, USA). Ions were analysed using a Top-10 data-dependent switching mode with the 10 most intense ions selected for HCD for peptide identification and reporter ion fragmentation in the Orbitrap. Automatic gain control targets were 30,000 for the iontrap and 1,000,000 for the orbitrap

### Quantitative MS Data analysis

Tandem mass spectra were extracted from the Xcalibur data system (version 2.2, ThermoScientific, USA) and searched through Mascot (v. 2.6.0) using Proteome Discoverer software (version 1.4.0.288, ThermoScientific, USA) to determine specific peptides and proteins. The parameters included: 20 ppm peptide precursor mass tolerance; 0.5 Da for the fragment mass tolerance; 2 missed cleavages, trypsin enzyme; TMT-6plex (N-terminus and K), carbamidomethyl (C) and oxidation (M) dynamic modifications; database: UniProt_HUMAN (release-2018_02, 20 366 entries). False discovery rate was set at 0.05 and 0.01 for relaxed and strict parameters respectively, with validation based on q-Value. The data were analysed using KNIME and embedded R scripts (KNIME analytics platform, Germany). Peptides were excluded from analysis if they were unassigned or had missing TMT channel intensity data; the primary accession number was taken for each peptide and proteins were grouped by this accession number with the geomean of individual peptide intensities given as the protein intensity value; TMT intensities were normalised using a sum scaling method and to the geomean of the two standard values for each peptide. Batches were then concatenated, batch corrected using ComBat (43) and PCA, clustering (XMeans and k-Means), gene ontology (GO) and specific protein analyses (fold changes and TTests) were carried out. Venn diagrams were produced using Venny 2.1 (http://bioinfogp.cnb.csic.es/tools/venny/). As the ComBat algorithm is only applicable to proteins present in all batches, a novel method of comparing samples across batches was developed. PCA plots of each non-ComBat corrected batch were carried out separately and Euclidean distances between each post-mouth rinse sample and the relevant unstimulated pool calculated. These Euclidean distances were then expressed relative to the distance between the two unstimulated pools which are present in each batch and, in theory, will vary to the same degree in each batch (Supplementary Figure a).

### Total protein concentration assay

The total protein concentration of collected saliva samples were determined by bicinchoninic acid assay (Thermo Scientific). Frozen saliva samples were defrosted on ice and then diluted 1:10 in ddH_2_0 in duplicate alongside a serial dilution of bovine serum albumin standard (2 mg/mL - 0.03125 mg/mL). Samples and standards were incubated with bicinchonic acid for 30 minutes prior to measuring absorbance as 540 nm using an iMark microplate absorbance reader (BioRad).

### Sodium dodecyl sulphate polyacrylamide gel electrophoresis

Sodium dodecyl sulphate polyacrylamide gel electrophoresis (SDS PAGE) was carried out on saliva samples. Saliva samples were prepared for electrophoresis by dilution 4x concentration LDS sample buffer (Invitrogen) with the addition of 0.5M DTT (Sigma) to the sample-buffer solution and then boiled for 3 minutes. Pre-cast 4-12% NuPage Novex Bis-Tris gels (Invitrogen) were assembled in a XCell vertical electrophoresis unit (Invitrogen) with MES running buffer (Invitrogen). Samples were loaded with equal protein concentration and electrophoresed for 32 minutes at 125 mA and 200 V (constant). Molecular masses were determined by comparison with SeeBlue Plus2 standard proteins (Thermo Scientific).

### Glycoprotein staining

Polyacrylamide gels were placed in 0.2% Coomassie Brilliant Blue R250 in 25% methanol and 10% acetic acid at room temperature for 90 minutes, followed by overnight de-staining in 10% acetic acid. Periodic acid Schiff’s (PAS) staining: 60 minute fixing in 25% methanol and 10% acetic acid, incubation with 1% periodic acid followed by water rinsing and Schiff’s reagent staining. Gels were imaged using the ChemiDoc MP Imaging System (BioRad).

### Immunoblotting

Separated proteins were electroblotted to nitrocellulose membranes for 60 minutes at 190 mA and 30 V (constant). Blots were blocked in 5% semi skimmed milk (Fluka) and probed with either an affinity-purified antibody fraction of mouse antiserum to a synthetic peptide of human cystatin-s corresponding to amino acid residues 21-141 (AF1296, R&D Systems) or an affinity-purified goat antibody raised against a peptide mapping at the C-terminus of human amylase (sc-12821, Santa Cruz). Binding was detected using a horseradish-peroxidase-labelled, affinity purified goat-ant-rabbit IgG (P0160, Agilent Dako) or rabbit-anti-mouse IgG (P0161, Agilent Dako) followed by Clarity Western ECL substrate detection system. Chemiluminescence was detected by ChemiDoc MP Imaging System (BioRad). Molecular masses were determined by comparison with SeeBlue Plus2 standard proteins (Thermo Scientific).

### Ethics

This study was approved by the King’s College London Ethics Committee (BDM/12/13-54).and written informed consent was obtained from all study participants.

### Statistical Analysis

Data were tested for normality using the Shapiro-Wilks normality test. 1-way ANOVA were used for determining statistically significant differences within the lower labial gland flow rates, parotid gland flow rates, protein output, cystatin S abundance datasets and, in the in-depth analysis, grouped WMS flow rate and protein output datasets. A 2-way ANOVA was used for determining statistically significant differences within the WMS flow rate datasets and, in the in-depth analysis, in the subject separated WMS flow rate and protein output datasets. The above analyses were carried out using Prism 6 software (GraphPad). The following were used to denote statistically significant differences in the figures: **** = P ≤ 0.0001, *** = P ≤ 0.001, ** = P ≤ 0.01, * = P ≤ 0.05.

### Data Availability

The PD 1.4 protein search file result containing accession numbers, percentage protein coverage, number of distinct peptides and quantification measurements can be found in Supplementary Tables 1-6. The raw-files and PD1.4 search files (protein and peptide) have been deposited to the ProteomeXchange Consortium via the PRIDE partner repository with the dataset identifier PXD017232 (Reviewer account details: Username; Password: o52lEXbo).

## Results

### TRP agonists stimulate salivary secretion

Significantly greater relative WMS flow rates were observed in response to the TRP agonist containing mouth rinses when compared to the UWMS flow rate (Figure 1a). Furthermore, 1 ppm nonivamide and 500 ppm menthol mouth rinsing significantly increased relative mean WMS flow rates compared to PG mouth rinsing, which itself significantly increased WMS flow rates compared to UWMS. The reproducibility of WMS flow rates in response to menthol and nonivamide mouth rinsing was demonstrated by repeating measurements with two of the participants (Figure 2a). All the mouth rinses increased mean WMS flow rate compared to unstimulated WMS (UWMS) flow rate (1.0 g/min). The highest concentrations of the three TRP channel agonists stimulated the greatest flow rates; 1.70 ml/min with 500 ppm menthol, 1.61 g/min with 300 ppm cinnamaldehyde and 1.67 g/min with 1 ppm nonivamide (Figure 2a (top)). When individual participants were considered, Figure 2a (bottom), we found that only participant 1 showed significantly greater stimulated flow rates.

**Figure 1.**
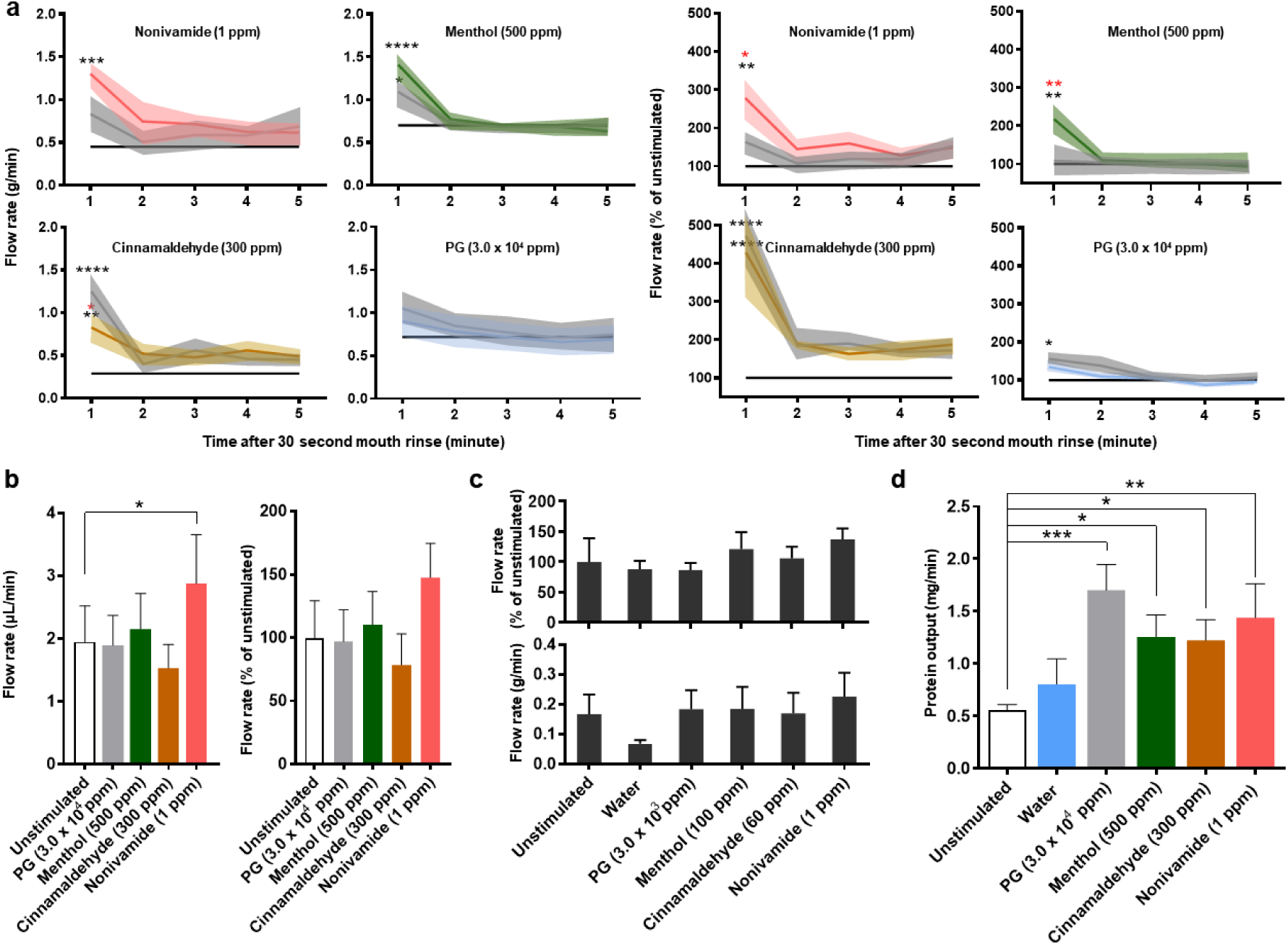
Effect of TRP channel agonists on salivary flow rates and protein output. a) WMS flow rate after 30 seconds of mouth rinsing expressed as absolute values (left) and relative to the unstimulated flow rate (right) (*n* = 6). Solid coloured lines indicate means and shaded areas indicate SEM. Grey indicates vehicle control (PG) at concentration used for the TRP agonist mouth rinse. Black line indicates mean unstimulated WMS flow rate. The blue line in the PG plots indicate water. Black * indicates significance versus unstimulated and red * indicates significance versus PG. b) Lower labial minor salivary gland flow rate after two minutes of mouth rinsing (Mean ± SEM; *n* = 10). c) Parotid saliva flow rate during two minutes of mouth rinsing (Mean ± SEM; *n* = 8). d) WMS protein output after 30 seconds of mouth rinsing (Mean ± SEM; *n* = 6).

**Figure 2.**
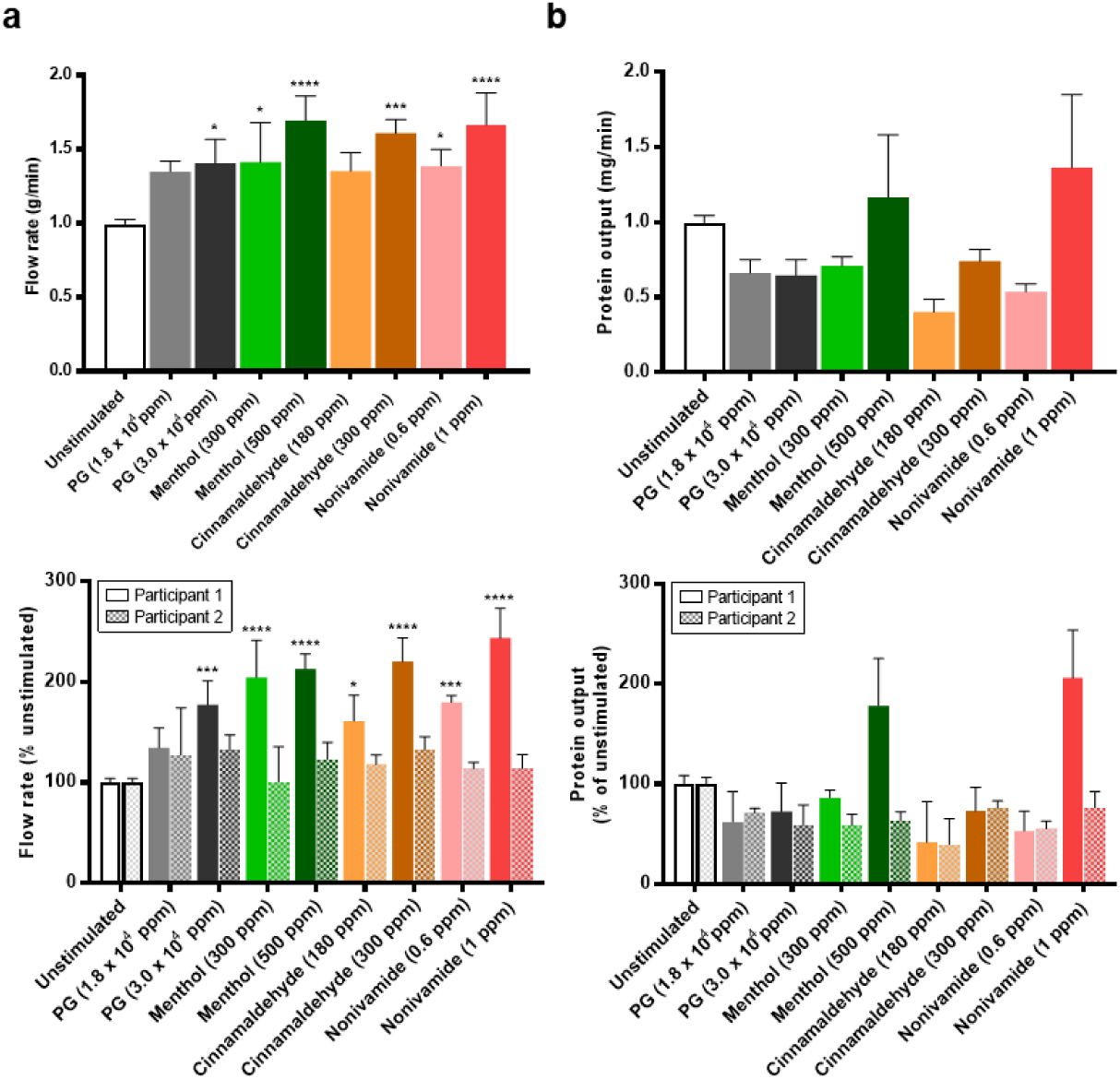
Reproducibility of the sialogogic properties of TRP channel mouth rinses. a) WMS flow rates of unstimulated saliva and stimulated saliva during the first minute after mouth rinse stimulation (top) and participant separated values relative to the unstimulated flow rate on the day of sampling (bottom). b) WMS protein output of unstimulated saliva and post-mouth rinse salivas in the two minutes after stimulation (top) and participant separated values relative to unstimulated protein output (bottom). All figures show mean ±SEM. Top figures: *n* = 6, unstimulated *n* = 48; Bottom figures: *n* = 3, unstimulated *n* = 24; *, **, *** and **** = P value from unstimulated ≤ 0.05, 0.01, 0.001 and 0.0001 respectively.

Nonivamide (1 ppm) mouth rinsing stimulated lower labial minor gland flow rate compared to the unstimulated flow rate (Figure 1b) but no mouth rinse caused parotid gland flow rates to significantly differ from unstimulated or water stimulated flows (Figure 1c).

TRP agonist mouth rinsing, as well as PG, caused greater WMS protein output (Figure 1d). These effects were shown to be less reproducible than the effects on flow rate (Figure 2b vs 3a). Although mean output in response to 1 ppm nonivamide (1.36 mg/min) and 500 ppm menthol (1.17 mg/min) were greater than UWMS (0.99 mg/min), these increases were not significant and can be attributed to participant 1, who showed a significantly greater response than participant 2 (Figure 2d).

#### Salivary proteomics overview

Overall 459 unique proteins were identified in saliva samples. The number of unique proteins identified in each of the 6 separate batches of samples varied from 199 to 158. Sixty four unique proteins were identified in all 6 sample batches (Figure 3a). Two reference proteomes were used to compare the proteins identified in this study to those identified in the literature. In a meta-analysis of proteins identified across six studies, Sivadasan et al. produced the largest publicly available “human salivary proteome”, consisting of 3449 unique human proteins (44). A second reference proteome was obtained from ProteomeDB (https://www.proteomicsdb.org/) which contained 1993 unique human proteins.

**Figure 3.**
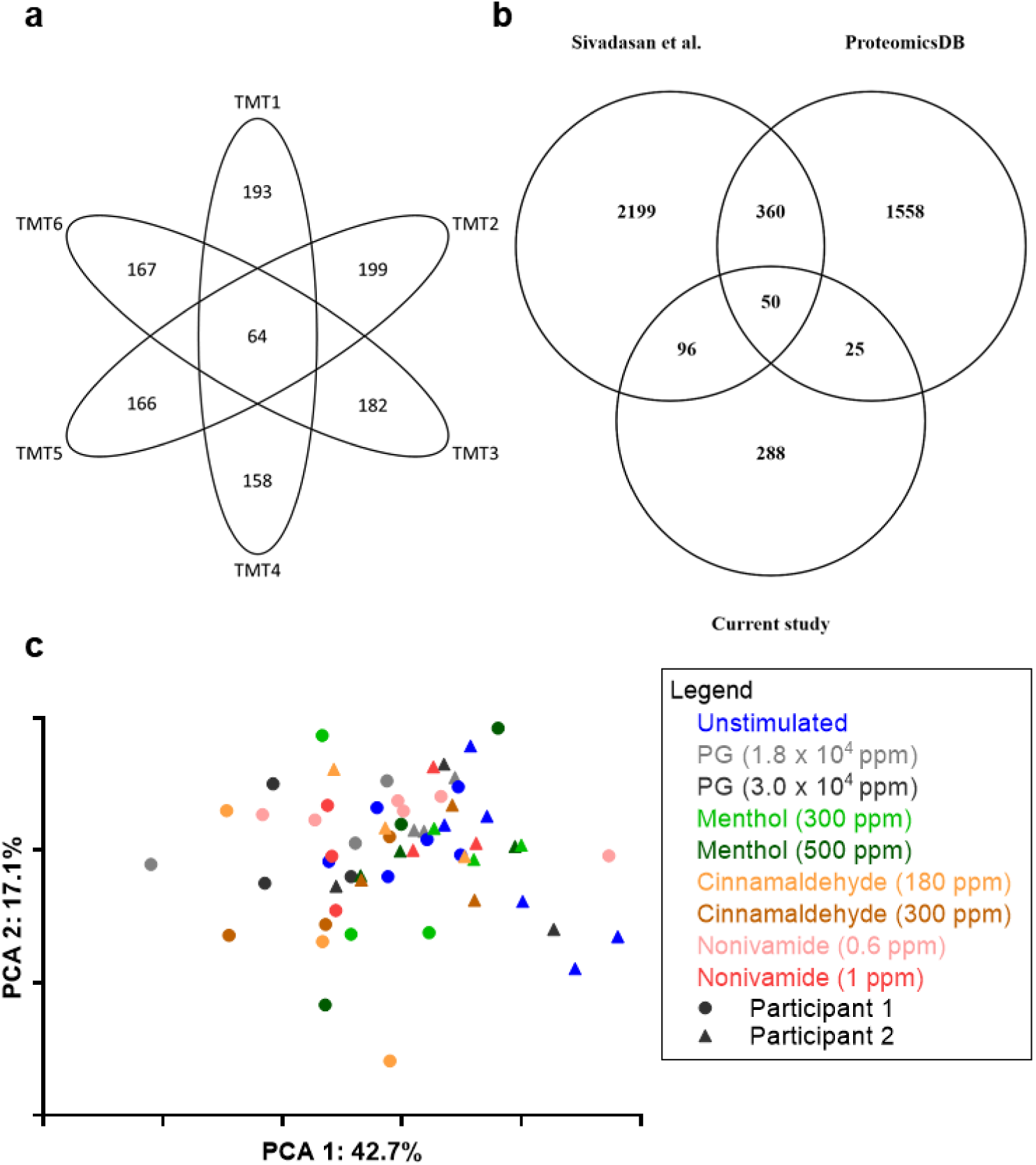
Proteomics overview. a) Venn diagram showing total number of identified proteins in each TMT10plex (outer) and the number of proteins identified in all TMT10plexes (inner) for all samples in each TMT10plex. b) Venn diagram showing the unique and common proteins identified in the current study, from a reference database (ProteomicsDB) and a meta-analysis of the salivary proteome by Sivadasan et al. 2015. c) PCA plot showing the distribution of unstimulated pools and post-mouth rinse WMS sample after ComBat batch correction.

Our study identified 288 unique human proteins absent from both datasets and so, to the best of our knowledge, are novel findings for the salivary proteome (Figure 3b). Greater confidence can be assigned to the 134 proteins that have a SwissProt annotation score of 5, relating to strong evidence of their existence *in vivo*, and of these, 12 were identified with at least one unique peptide across the batches, of which 9 had a relative abundance of less than 0.2%.

### Sources of variation in the salivary proteome

When all samples were labelled by participant and condition (Figure 3c), it is clear that samples are discriminated by participant along the x-axis (PCA1). Furthermore, if the geomean of the replicates of each condition are taken (Figure 4) and k-means clustering (number of clusters having been determined by x-means) applied then 100% of participant 2 samples cluster together and 89% of participant 1 samples cluster together. All stimulated samples from participant 2 clustered separately from the unstimulated sample, reflecting that this subject was a responder. In contrast none of the stimulated samples from participant 1 clustered separately from unstimulated samples, reflecting that this subject was a non-responder. Since the x-axis represents the principal component responsible for the majority of the variation in the dataset (57.1%), we conclude that the person the saliva comes is the major source of variation between WMS proteomes.

**Figure 4.**
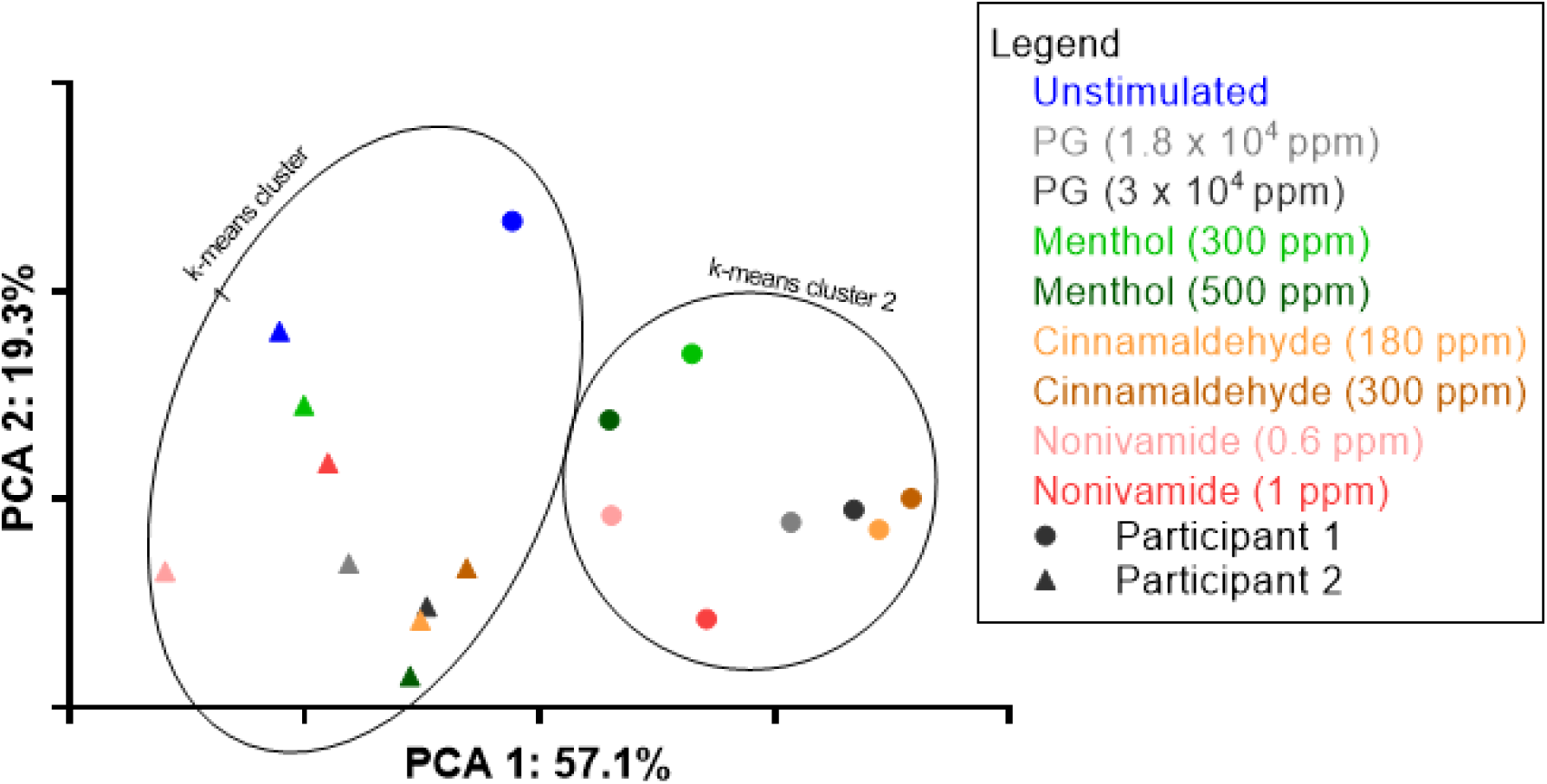
Identification of sources of variation in the salivary proteome. a) PCA plot showing the distribution of the geomean of each of the sample conditions with highlighted k-means clusters

The geomeans of post-mouth rinse samples were separated by mouth rinse primarily on the y-axis of Figure 4, representing the principal component responsible for 19.3% of variation in the dataset. For both participants, post-PG and cinnamaldehyde mouth rinse coordinates associated together, suggesting that the cinnamaldehyde mouth rinses were not causing additional variation in the WMS proteome than was already induced by the PG in the mouth rinse. However, post-nonivamide and menthol coordinates were separated from the PG coordinates suggesting these compounds were inducing proteome changes independently of PG (note the lower concentrations of PG in nonivamide and menthol mouth rinses compared to cinnamaldehyde (Table 2).

Supplementary Figure b shows the mean (±SEM) variability of each post-mouth rinse sample to the unstimulated pool in both participants. Nonivamide caused changes in the WMS proteome in both participants, 1 ppm in participant 1 and 0.6 ppm in participant 2. Cinnamaldehyde (300 ppm) and to a lesser degree menthol (300 ppm) caused relatively large changes in the WMS proteome of participant 1. Large variation was sometimes seen in the proteome response to the same mouth rinse in the same participant, as indicated by the large SEM values, for example in participant 1-300 ppm menthol and participant 2-0.6 ppm nonivamide. In contrast, some mouth rinses cause very repeatable changes, for example 300 ppm menthol in participant 2 and 0.6 ppm nonivamide in participant 1.

### Specific protein changes

Ten unique proteins were significantly regulated by TRP channel agonist stimulation (Table 4), five of which belong to the cystatin family. Salivary cystatins (S, SA or SN) were upregulated in response to every mouth rinse with the greatest degree of upregulation observed in response to nonivamide mouth rinses. The peptides assigned to each of these proteins (13, 10 and 17 to S, SA and SN respectively) were unique. Additionally, cystatin D was upregulated at both concentrations of nonivamide and cystatin C was upregulated after 1 ppm nonivamide mouth rinsing. Menthol at 500 ppm caused upregulation in salivary cystatins to a greater extent than PG. Although salivary cystatins were upregulated after cinnamaldehyde mouth rinsing, it was less than with PG mouth rinses despite the same concentration of PG being present in 1.8 × 10^4^ ppm and 3.0 × 10^4^ ppm PG to 180 ppm and 300 ppm cinnamaldehyde respectively. The finding that salivary cystatins are upregulated by 1 ppm nonivamide mouth rinsing was supported by qualitative immunoprobing (Figure 5). Statistically significant greater cystatin S was observed in WMS after 1 ppm nonivamide mouth rinsing (Figure 5c).

**Table 4.** WMS proteins regulated by TRP channel agonist mouth rinsing Fold change in geomean (compared to unstimulated saliva) of WMS proteins after rinsing with TRP channel agonist or vehicle with significant regulation (p < 0.05) across both participants. Fold changes recognised as up- or downregulated are highlighted in green and red respectively. Blanks indicate that protein was present but not regulated. Additionally: the total number of peptides identified across all 6 batches is reported as well as the mean protein coverage across the six batches.

| Protein ID | Protein Name | Total peptides identified (% of total) | Mean protein coverage (%) | PG |  | Cinnamaldehyde |  | Menthol |  | Nonivamide |  |
| --- | --- | --- | --- | --- | --- | --- | --- | --- | --- | --- | --- |
|  |  |  |  | 1.8 x 10<sup>4</sup> ppm | 3.0 x 10<sup>4</sup> ppm | 180 ppm | 300 ppm | 300 ppm | 500 ppm | 0.6 ppm | 1 ppm |
| P12273 | Prolactin-inducible protein | 258 (0.95) | 13.58 | 1.92 | 1.82 | 1.60 | 1.73 |  |  |  |  |
| P59665 | Neutrophil defensin 1 | 367 (1.35) | 24.83 | 1.62 | 1.57 |  |  |  |  |  |  |
| P01034 | Cystatin-C | 205 (0.76) | 40.41 |  |  |  |  |  |  |  | 1.56 |
| P28325 | Cystatin-D | 202 (0.75) | 31.80 |  |  |  |  |  |  | 1.64 | 1.79 |
| P01036 | Cystatin-S | 1227 (4.53) | 76.59 |  | 1.57 | 1.59 | 1.61 |  | 1.66 | 1.81 | 1.72 |
| P09228 | Cystatin-SA | 326 (1.2) | 38.89 |  | 2.08 | 1.72 | 1.87 | 1.77 | 2.02 | 2.15 | 2.14 |
| P01037 | Cystatin-SN | 4024 (14.84) | 66.55 |  | 1.52 |  |  |  | 1.68 | 1.82 | 1.79 |
| P01860 | Ig gamma-3 chain C region | 74 (0.27) | 14.15 |  |  | 0.52 |  |  |  | 0.63 | 0.56 |
| Q9BWT7 | CARD10 | 74 (0.27) | 1.45 |  |  | 0.66 | 0.66 |  |  |  |  |
| P00558 | Phosphoglycerate kinase 1 | 60 (0.22) | 7.00 |  |  | 0.43 |  |  |  |  |  |

**Figure 5.**
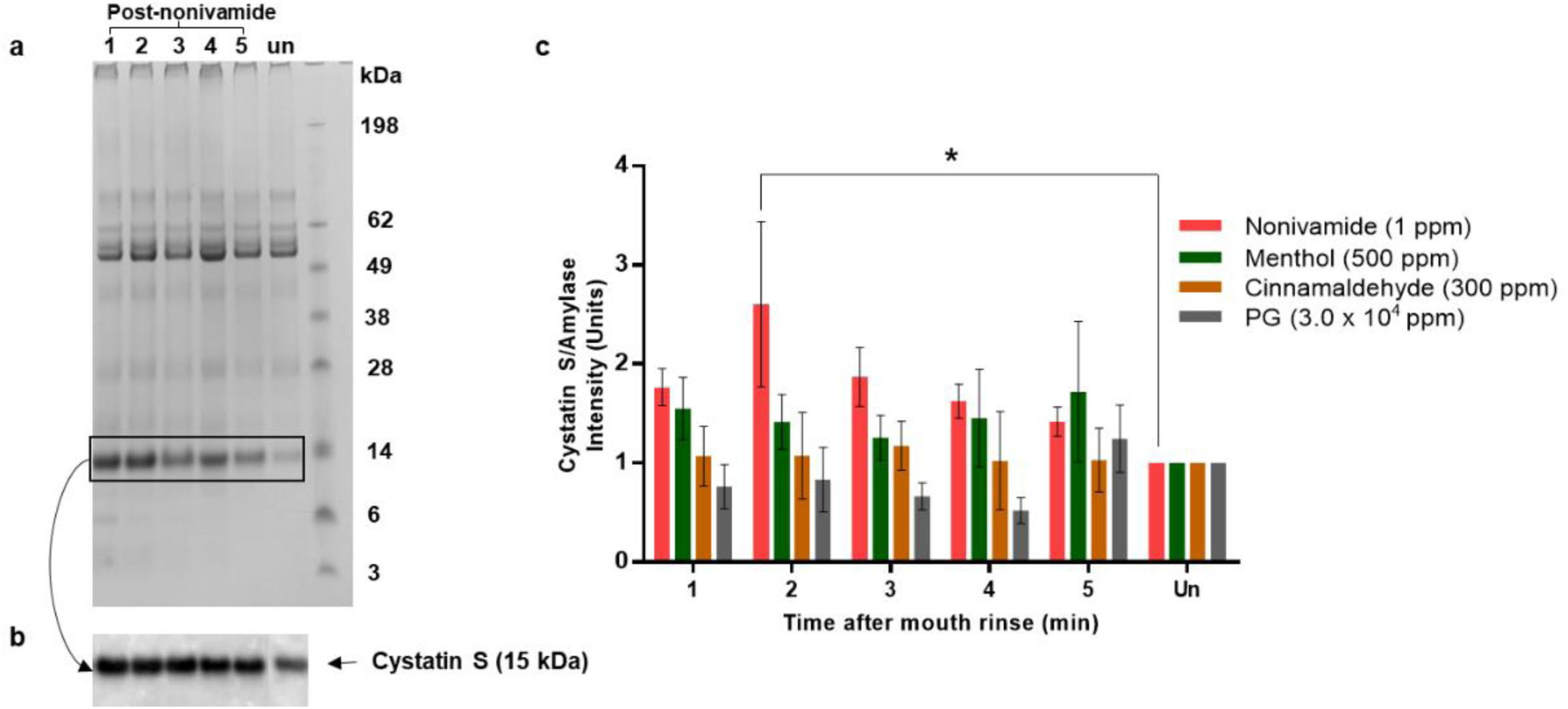
WMS cystatin S abundance after TRP channel agonist mouth rinsing. a) An example of Coomassie blue and PAS stained salivary proteins separated by SDS PAGE from one participant demonstrating how the cystatin S band intensities increase after nonivamide b) Western blot of the same samples as in a) identifying the protein band as cystatin S. (un: unstimulated, 1 - 5: 1 - 5 min after mouth rinse. c) Intensity of the cystatin S band on a western blot, relative to the amylase western blot band intensity, in WMS collected after a 30 second TRP agonist mouth rinse normalised to unstimulated saliva (Mean±SEM; n = 6).

Two other proteins were upregulated in the dataset, prolactin-inducible protein was upregulated after both PG and cinnamaldehyde mouth rinsing whilst neutrophil defensin 1 (α-defensin) was upregulated in response to PG (Table 4). Cinnamaldehyde (180 ppm) resulted in the down-regulation of IgG-3 chain C region, caspase recruitment domain-containing protein 10 (CARD10) (also downregulated in 300 ppm cinnamaldehyde) and phosphoglycerate kinase 1 (PGK1). IgG-3 chain C region was also downregulated in response to nonivamide.

## Discussion

In this study we have found that mouth rinsing with menthol or nonivamide increases WMS flow rate (Figure 1 & Figure 2). These observations expand on the current reports in the literature that TRPV1 agonists, such as piperine, nonivamide, capsaicin and 6-gingerol can stimulate salivary secretion since stimulation of salivary secretion by menthol has not previously been described. We have further found that nonivamide can stimulate minor gland secretion. Cinnamaldehyde mouth rinse did not evoke a salivary response even though it was perceived to be as intense or more intense than the menthol or nonivamide mouth rinses (Supplementary data c), which indicates that salivary responses are TRP agonist specific. The effect of a cinnamaldehyde mouth rinse was no greater than the vehicle PG but both were greater than unstimulated WMS (Figure 1a). Nonivamide, menthol and PG increased outputs of total protein in saliva suggesting that the protein composition and properties of saliva might be altered. Cinnamaldehyde decreased protein secretion compared to the PG vehicle. This is likely due to cinnamaldehyde diminishing the sialogogic properties of PG through a reaction between the compounds rather than inhibiting the nerve mediated reflex PG induces as no inhibitory neurones exist (45). The source of increased protein secretion is presumably salivary gland exocytosis of protein storage granules but it may be that there are other contributions from within the oral cavity. In order to investigate further, quantitative changes in salivary protein composition we implemented a bottom-up mass spectrometry pipeline new to salivary proteomics, which led to the identification of novel whole WMS proteome changes and specific protein changes in response to the TRP channel agonists studied. From PCA we identified that the largest source of variation in the salivary proteome was between subjects but that changes in the proteome were also caused by different mouth rinses (Figure 4). Repeat analyses on subjects demonstrated that there was variation from day to day in response to some of the mouth rinses.

The mass spectrometry pipeline applied in this study produced results that contribute to the salivary proteome literature, since it identified proteins in saliva that have not previously been reported (Supplementary Table). This may be due to the novel application of IEF using OFFGEL electrophoresis with TMT labelled quantitative tandem mass spectrometry LC-MS/MS to salivary proteomics but may also be the result of searching against updated databases or inter-personal differences in salivary composition, which has previously been observed to have a larger coefficient of variation than intra-personal variation (46). Three previous studies of WMS have used IEF in tandem mass spectrometry (47-49), and a further study coupled it with mTRAQ quantification methodology (50). However, these studies did not couple IEF with isobaric labelling such as TMT. It could be that the novel methodology contributes to better identification of lower abundance proteins, or this could be a result of the experimental stochasticity in bottom-up mass spectrometry approaches, the use of updated protein sequence database or differences in raw data analysis software. Despite being in lower abundance, the novel proteins are of sufficient length (median amino acid length being 897 and ranging from 97 to 7570) to produce detectable tryptic peptides. This suggests that the method is not just identifying small proteins with a high abundance but proteins of a range of sizes with relative abundances ranging from 3.2% of total peptides to < 0.005% (Supplementary table). A bottom-up approach was implemented with the intention to maximise the quantification of the salivary proteome. With 459 proteins quantified, the coverage was limited when compared to other TMT quantification studies with more state of the art equipment. Furthermore, good proteome coverage that also represents the variety of gene products has been achieved in top-down and data independent acquisition proteomic studies and could be used to further investigate the diversity of the salivary proteome (51, 52).

The presence of some lower abundance proteins appeared to be influenced by mouth rinsing, for example CARD10 and phosphoglycerate kinase 1 (PGK1), which were 0.3 and 0.2% of total identified peptides respectively (Table 4). This is the first time CARD10 has been identified in WMS. Both CARD10 and PGK1 were downregulated specifically in response to cinnamaldehyde mouth rinsing:. Despite there being no previous reports of association between CARD10 and cinnamaldehyde, there have been previous reports of cinnamaldehyde inhibiting other caspase recruitment domain proteins in mice and subsequent anti-inflammatory effects (10). Similarly, there have been no previous reports of an association between cinnamaldehyde and PGK1. However, anti-angiogenesis properties of cinnamaldehyde and cinnamon extract have been previously reported (12-14). The observation of down-regulation of CARD10 and PGK1 could be preliminary evidence that the anti-inflammatory and bactericidal effects of cinnamaldehyde extend to short term mouth rinsing in the oral cavity.

Upregulation of cystatin S in the WMS secreted in response to nonivamide was detected by mass spectrometry and western blotting (Figure 5). Despite significant sequence homology between the salivary cystatins, the peptides assigned to S, SN and SA were unique to each protein. Furthermore, the antibody used in western blotting had a reasonable specificity for cystatin S, with 30% and 5% cross reactivity to cystatins SN/SA or D/C respectively. To further increase the confidence in specificity, a top down approach could be used as demonstrated in the literature (53). Greater quantities of cystatin S in saliva could result in an improvement in mucosal adhesion, a property of saliva important in mouthfeel and xerostomia. Cystatin S has been shown to interact with oral mucosal surfaces and play a role in the formation of protein pellicles *in vitro* on hydrophobic surfaces that mimic the mucosa (54). Coupled with previous observations that the rheological properties of saliva are modified by nonivamide (29, 55), mouth rinsing with nonivamide as a treatment for xerostomia warrants further study. Increased cystatin S expression may have other potential benefits for oral health. due to inhibition of cysteine protease activity, as indicated by significant enrichment of the “negative regulation of cysteine-type endopeptidase activity” GO. The upregulation of the GO for cysteine protease inhibition mirrors the western blotting findings and work in the literature (56, 57). Cystatin S has been shown to inhibit proteolytic activity in the culture supernatant of *P. gingivalis* (58), a Gram negative bacterial species that produces the gingipain class of cysteine proteases which are implicated in periodontal disease (59). Additionally, cystatin S, as well as prolactin-inducible protein, upregulation could improve acceptance of bitter taste as indicated by the GO enrichment “detection of chemical stimulus involved in sensory perception of bitter taste” (60). This suggests that TRPV1 agonists could be used to promote the consumption of bitter foods, the reduced consumption of which has been implicated in the health, dietary intake and weight of “super tasters” (61).

This study is the first to demonstrate an acute salivary cystatin S response to TRPV1 agonists in humans (Figure 5). A cystatin S-like protein response to capsaicin has been demonstrated in rats fed on a capsaicin-adulterated diet; the presence of a new protein in rat saliva was demonstrated and the protein found to have cystatin S-like properties such as inhibition of cysteine protease activity (57). In the rat increased cystatin S-like protein levels enhanced consumption of a capsaicin rich diet and it was hypothesised that this response may be triggered by irritation of the oral mucosa (56). Although these studies, along with the current study, both show increases in cystatin S and cystatin S-like proteins in saliva, the time scales over which the phenomenon occurs are significantly different. The current study shows the reversible increase within two minutes of nonivamide mouth rinsing whilst in the studies in rat the increase was observed after three days of capsaicin-adulterated diet, suggesting different mechanisms are responsible. The increase in cystatin S levels in WMS in the current study must be due to the release of preformed protein as it takes 30 minutes for newly synthesised protein containing vesicles to pass from the rough endoplasmic reticulum to the condensing vacuoles in secretory cells (62).

The identification of proteins regulated across all mouth rinses alongside proteins only regulated in response to one mouth rinse suggests, in agreement with the total protein secretion data, that there are different mechanisms responsible for the regulation of proteins in WMS. Furthermore, some of the proteins are known to be produced by the salivary glands whereas others are non-salivary proteins. The upregulation of salivary cystatins (S, SN and SA) may reflect a preferential stimulation of the submandibular/sublingual glands, the primary producers of salivary cystatins (63). Cystatin S regulation may be influenced by direct effects of the agonists on minor glands, as lower labial gland flow rates were greater after 1 ppm nonivamide mouth rinsing (Figure 1b) and they have been demonstrated to express cystatin S and other salivary proteins (64). Menthol, cinnamaldehyde and nonivamide are highly lipophilic compounds, having partition coefficient values (an indicator of lipophilicity; higher values imply greater lipophilicity) of 3, 1,9 and 4.2 respectively. Comparatively, pilocarpine, a drug that has previously been used to directly stimulate minor salivary glands (65), has a partition coefficient value of 1.1 (66). Higher lipophilicity suggests that these TRP channel agonists would have a greater permeability in the oral epithelium and lamina propria than pilocarpine, which would enhance direct activation of TRP channels expressed in minor glands.

The significantly greater WMS flow rates observed in the proteomics study (Figure 2a) are primarily the result of the response from one of the two participants, with the other showing little response to the TRP agonists. There is a precedence in sensory science for responders/non-responders, such as in the case of the detection of the bitter compound PROP which is associated with the expression of the TAS2R28 bitter receptor gene (67). Although the comparison seems to be limited by the fact that participants in the current study do have a sensory perception of the TRP agonists, the mechanism for salivary secretion in response to TRP agonist detection is yet to be elucidated and unknown genetic factors could be responsible for the prevalence of salivary non-responders to TRP agonists despite a sensory perception. A breakdown of the dataset shown in Figure 1a reveals that only 2 of the 19 participants given a TRP containing mouth rinse did not exhibit an increase in WMS flow rate (as defined by a flow rate 150% that of unstimulated flow rate). This suggests that the prevalence of non-responders in the population is lower than the 50% suggested in the proteomics study.

In summary this study provides the first evidence for stimulation of salivary secretion by a non-TRPV1 TRP channel agonist. Increased minor gland secretion may be a direct action of the TRP agonists on submucosal salivary glands alongside nerve-mediated mechanisms. Furthermore, novel changes in the proteome of the saliva secreted in response to the TRPV1 agonist nonivamide were identified by mass spectrometry and supported by western blotting. These findings suggest that TRP channel agonists can be explored as potential candidates for altering salivary secretion, particularly in subjects with xerostomia and reduced levels of saliva.

## Abbreviations

CARD10: CAspase Recruitment Domain-containing protein 10
DTT: DiThioThreitol
LC-MS/MS: Liquid Chromatography - tandem Mass Spectrometry
GO: Gene Ontology
IEF: IsoElectric Focusing
LDS: Lithium Dodecyl Sulphate
MS: Mass Spectrometry
PG: Propylene Glycol
PGK1: PhosphoGlycerate Kinase 1
TCEP: Tris (2-CarboxyEthyl) Phosphine
TEAB: TriEthylAmmonium Bicarbonate
TMT: Tandem Mass Tag
TRP: Transient Receptor Potential
TRPA1: Transient Receptor Potential cation channel, subfamily A, member 1
TRPM8: Transient Receptor Potential cation channel subfamily M member 8
TRPV1: Transient Receptor Potential cation channel subfamily V member 1
WMS: Whole Mouth Saliva
UWMS: Unstimulated Whole Mouth Saliva

## Acknowledgements

The authors would like to acknowledge a BBSRC PhD Case Studentship subsided by Symrise AG as the source of funding for the work. Grant number: BB/L015498/1.

## Data Availability

The raw-files and PD1.4 search files (protein and peptide) have been deposited to the ProteomeXchange Consortium via the PRIDE partner repository with the dataset identifier PXD017232 (Reviewer account details: Username; Password: o52lEXbo).

